# Increased population density depresses activity but does not influence emigration in the snail *Pomatias elegans*

**DOI:** 10.1101/2020.02.28.970160

**Authors:** Maxime Dahirel, Loïc Menut, Armelle Ansart

**Affiliations:** Univ Rennes, CNRS, ECOBIO (Ecosystèmes, biodiversité, évolution) – UMR 6553, F-35000 Rennes, France; INRAE, Université Côte d’Azur, CNRS, ISA, France

**Keywords:** Defence, dispersal syndromes, ecosystem functioning, gastropod, sex-biased dispersal

## Abstract

Dispersal is a key trait linking ecological and evolutionary dynamics, allowing organisms to optimize fitness expectations in spatially and temporally heterogeneous environments. Some organisms can either actively disperse or reduce activity in response to challenging conditions, and both responses may be under a trade-off. To understand how such organisms respond to changes in environmental conditions, we studied emigration (the first step of dispersal) and activity behaviour in the gonochoric land snail *Pomatias elegans*, a litter decomposer that can reach very high local densities, over most of the range of ecologically relevant densities. We found that crowding had no detectable effect on emigration tendency in this species, contrary to previous results in many hermaphroditic snails. *Pomatias elegans* is nonetheless able to detect population density; we show they reduce activity rather than increase dispersal in response to crowding. We propose that limiting activity may be more advantageous than moving away in species with especially poor movement abilities, even by land mollusc standards, like *P. elegans*. Interestingly, emigration and activity were positively correlated independently of density; this dispersal syndrome may reflect an underlying pace-of-life syndrome, and is compatible with a dispersal-dormancy trade-off, which would require further investigation. Additionally, we found snails with heavier shells relative to their size tended to be less mobile, which may reflect physical and metabolic constraints on movement and/or survival during inactivity. We finally discuss how the absence of density-dependent dispersal may explain why *P. elegans* is often found at very high local densities, and the possible consequences of this behaviour for ecosystem functioning and litter decomposition.

## Introduction

Dispersal, i.e. movement potentially leading to gene flow in space, is a key organismal trait influencing and linking ecological and evolutionary dynamics in sometimes complex ways (Ronce, 2007; Clobert *et al*., 2012; Bonte & Dahirel, 2017). As the cost-benefit balance of dispersal is expected to vary depending on individual phenotype and experienced conditions, dispersal is often highly context- and phenotype-dependent (Bowler & Benton, 2005; Clobert *et al*., 2012; Baines, Ferzoco & McCauley, 2019; Endriss *et al*., 2019). Complexity is even added when one considers that dispersal can be seen as a three-step process (emigration, transfer, immigration), and that different factors can have different levels of influence at each step (Matthysen, 2012). It has been hypothesized (e.g. Matthysen, 2012; Bonte & Dahirel, 2017), that this multicausality of dispersal is why even well-studied environmental factors may lead to very different dispersal responses depending on populations and species. To understand the context-dependency of dispersal, it is thus necessary to study a broader range of life histories and contexts, including organisms that may have alternative strategies to escape negative environmental conditions, e.g. dormancy (Buoro & Carlson, 2014).

The effects of population density on dispersal, and especially emigration, are likely among the most studied of all dispersal drivers (Bowler & Benton, 2005; Matthysen, 2005; Harman *et al*., 2020). However, while emigration increasing with density and competition would be a straightforward expectation, and is supported by theory (Travis, Murrell & Dytham, 1999; Rodrigues & Johnstone, 2014), empirical results are actually much more idiosyncratic both within and among species: positive density-dependent, negative density-dependent and density-independent emigration seem to occur at broadly similar frequencies (Jacob *et al*., 2019; Harman *et al*., 2020). Explanations for this variability typically invoke multicausality and interactions with other traits, such as sociality (with more social species/individuals in a species less likely to leave large groups, leading to negative density-dependent dispersal⍰; Serrano *et al*., 2003; Cote & Clobert, 2007), body size (as larger and smaller individuals do not feel competition with the same intensity; e.g. Baines *et al*., 2019), or sex and sex-ratio (as competition for resources and for partners interact; De Meester & Bonte, 2010; Hovestadt, Mitesser & Poethke, 2014), among many others. Despite the large amount of studies, we are still far from integrating them into a general understanding of density-dependent dispersal: a possible reason is that invertebrates and especially non-arthropod invertebrates are under-represented in the published literature (Harman *et al*., 2020), meaning we have only explored a narrow part of the trait combinations that may interact with, and shape, the density-dependence of dispersal.

The land winkle *Pomatias elegans* (Müller 1774)(**Fig. 1A**), is a common gastropod species in forests and hedgerows across Europe (Kerney & Cameron, 1999; Falkner *et al*., 2001; Welter-Schultes, 2012). As a caenogastropod, it has separate sexes, contrary to the majority of land molluscs which are hermaphrodite (Barker, 2001). Its active dispersal capacities are poorly studied but expected to be very low, even by molluscan standards (Kramarenko, 2014); average dispersal distances after 42 days are below 20 cm, according to one of the only (indirect) records available in the peer-reviewed literature (Pfenninger, 2002). Like many other land snails (Cook, 2001; Falkner *et al*., 2001), it is able to survive up to several months under unfavourable conditions by strongly reducing its activity (sometimes down to dormancy; Kilian, 1951; Falkner *et al*., 2001). *P. elegans* is often distributed in a very aggregated way across landscapes, and when a population is present, local densities can vary by several orders of magnitude, from less than one to several hundred individuals.m^−2^ (Pfenninger, 2002). When it is present, *P. elegans* plays a key role in ecosystem functioning, by decomposing litter and altering its chemical and physical characteristics (Coulis *et al*., 2009; De Oliveira, Hättenschwiler & Handa, 2010).

**Figure 1.**
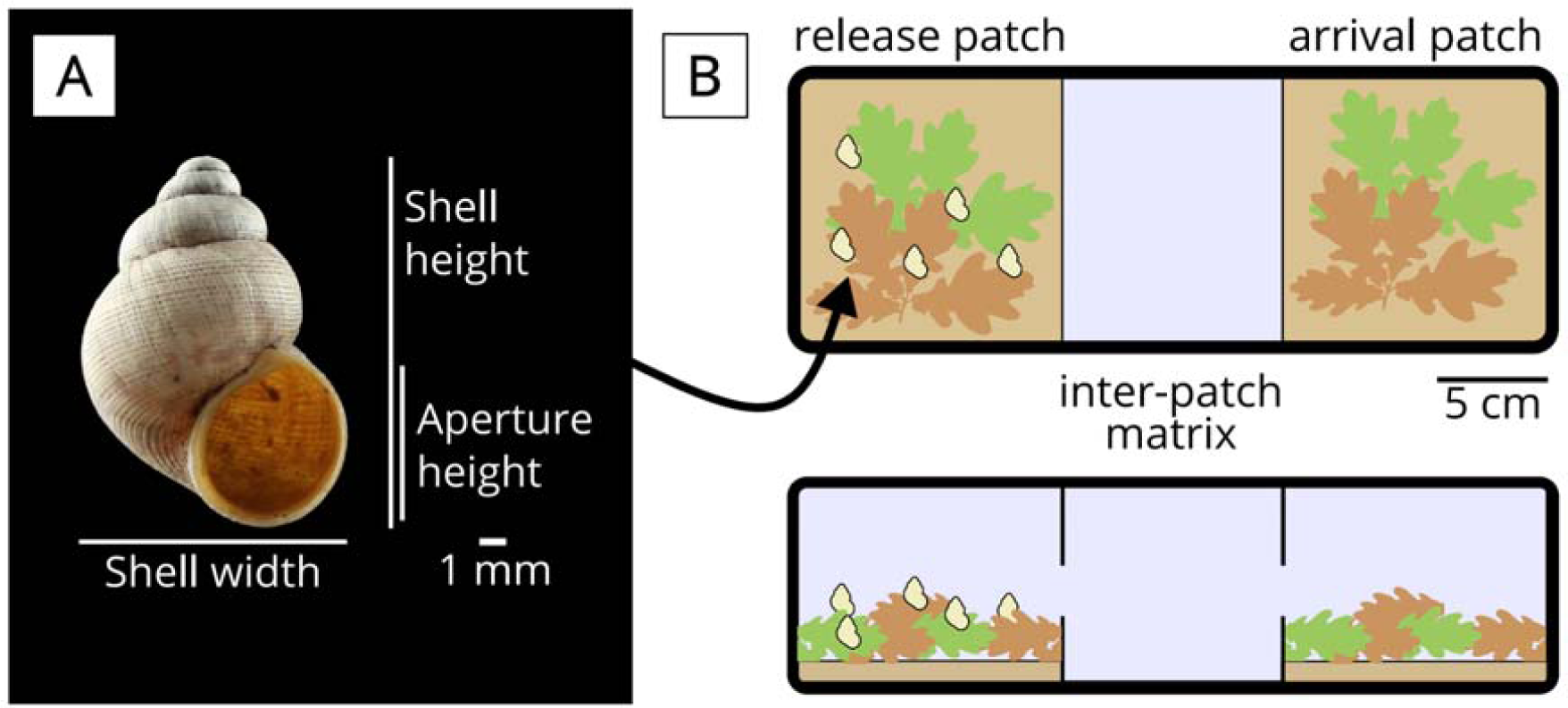
(A) *Pomatias elegans* shell showing the measurements used in the present study. (B) Diagram of the experimental two-patch system, showing (to scale) the release and arrival patches containing litter and soil, and the empty matrix in-between. Top: viewed from above; bottom: side view. Credits: snail photograph – Armelle Ansart; derived snail silhouette – Maxime Dahirel (deposited in phylopic.org); *Quercus robur* leaves silhouette – Ferran Sayol, available in phylopic.org under Public Domain Dedication 1.0 license.

We here tested the hypothesis that emigration (the first step of dispersal) and activity were density-dependent in *P. elegans*. We tested both activity and actual emigration for two reasons. We expected dispersers to be more active than residents, density being equal, in agreement with general predictions on behavioural syndromes (Cote *et al*., 2010; Réale *et al*., 2010). However, the two traits may not be interchangeable, and divergent responses between the two may help understand how *P. elegans* cope with high densities. For instance, if activity merely is a correlate of dispersal, or vice-versa (for instance if dispersal is just a “by-product” of routine movement; Van Dyck & Baguette, 2005), we would expect activity to respond to density in the same way as emigration. If reduced activity is also an alternative coping strategy in the face of stress, we may expect instead a depression of activity in response to density (Cameron & Carter, 1979). In accordance with the phenotypic compensation hypothesis, which predicts that risk-taking behaviour should be associated with morphological traits limiting vulnerability to predators (e.g. Ahlgren *et al*., 2015), we also expected dispersers/active individuals to be better defended, by having heavier shells. Finally, we accounted for sex differences, given sex-biased dispersal is often observed in other taxa with separated sexes (Trochet *et al*., 2016).

## Material and methods

We collected *Pomatias elegans* snails in spring 2017 in leaf litter and at the bottom of calcareous walls (Falkner *et al*., 2001) in Arcais, France (approximate location: 46°17’60”N, 0°41’21”W). They were then kept under controlled conditions (20°C, L:D 16:8) in plastic boxes (25 × 25 × 9 cm, about 500 snails.m^−2^) with a 1 cm layer of soil and a roughly 2-3 cm thick layer of leaf litter collected in urban and suburban woodlots and hedgerows on and close to the University of Rennes Beaulieu campus (predominantly oaks *Quercus robur* L. 1753 and *Q. petraea* (Matt.) Liebl. 1784, and beech *Fagus sylvatica* L. 1753). We provided humidity by watering soil and litter every 10 to 15 days.

### Dispersal experiment

In June 2017, we created two-patch systems by adapting the system we developed for slugs in Fronhofer et al. (2018), but with a shorter inter-patch gap and longer dispersal period to account for the more limited dispersal capacities of *P. elegans* (**Fig. 1B**). We divided 32 × 12 cm^2^ transparent plastic boxes (height: 9 cm) in two 12 × 12 cm^2^ patches (“release” and “arrival”) and one central 10 × 12 cm matrix, using plastified cardboard walls. A 2 × 12 cm^2^ slot was left open in each wall to allow snails to leave and enter patches. Both patches were filled with watered soil and litter as in maintenance boxes (see above), while the matrix was kept dry and empty. Experiments were done under controlled conditions (20°C, L:D 16:8).

We then placed between 1 and 30 snails in each “release” patch, and first closed the patches for 24 h to let the snails habituate. We then let them free to move for 4 days, and recorded which snails stayed in their release patch (hereafter “residents”) and which snails emigrated to the arrival patch. No snails were found in the matrix between the two patches, and no snails died during the experiment. We tested the following group sizes: 1, 2, 3, 4, 5, 10, 15, 20, and 30 snails per box, with respectively 30, 20, 10, 9, 6, 4, 3, 3, and 2 replicates (total: 371 snails in 87 replicates). We chose these replicate numbers to have at least 30 individuals and 2 replicates per group size. We focused our replication effort more on individuals than boxes (although we did both to some extent), as we were interested in individual traits both as responses and covariates (see Statistical analyses), so a substantial number of individuals at each density was needed. The selected group sizes correspond to initial densities ranging from 69.4 to 2083.3 snails.m^−2^ (taking only release patch surface area into account), and likely provide a good coverage of natural densities. Indeed, densities of up to 1072 detected snails.m^−2^ have been found using 25 × 25 cm^2^ quadrats in natural populations (Pfenninger, 2002), and a separate study indicates this species has a field detectability in the 0.5 to 0.9 range when hand collected (Albano *et al*., 2015), possibly because many individuals may have buried themselves in the substrate at any one time (Falkner *et al*., 2001). We aimed and managed to ensure balanced sex ratios across densities (in line with natural populations⍰; Boycott, 1916); see **Supplementary Material 1** for a description of the sexing procedure, including both (shell morphology-based) putative sexing before the experiment and (dissection-based) confirmatory sexing after. Tests were spread haphazardly over the month due to logistical constraints (start dates ranging from June 2 to June 26); no reproduction was observed during this interval.

### Post-dispersal phenotyping

Within 24h after dispersal tests, we placed each snail on a small (8 cm diameter) clean patch of wet substrate (synthetic upholstery foam) and recorded the latency to activity, snails being defined as active if their operculum was open and both their foot and tentacles were visible out of the shell. Experiments were done under controlled (20°C) temperature conditions, always during the “light” period. All snails that had not moved after 20 minutes remained inactive, operculum closed, for at least two hours, indicating they were not simply slower individuals, but actually behaved differently from the active ones. Indeed, a limited number was further monitored and remained fully inactive even after 7 hours. We therefore analysed activity as a binary variable (active within 20 minutes/ not active for at least two hours). After sexing by dissection (see **Supplementary Material 1**), shells were dried at room temperature before being weighed twice (to the nearest 0.1 mg; CP224S scale, Sartorius, Göttingen, Germany). Twenty-four out of 371 shells were partly broken during dissection and could not be weighed.

### Statistical analyses

All analyses were done in a Bayesian framework, using the probabilistic language Stan, which uses a variant of Hamiltonian Monte Carlo (NUTS) as an MCMC method to obtain satisfactory posterior samples in a much lower number of iterations than other methods (Carpenter *et al*., 2017; Stan Development Team, 2018). We created our Stan models using R (R Core Team, 2020) and the *brms* package (Bürkner, 2017) as front-ends; this allows models to be specified using R formula syntax. Scripting, analysis and plotting relied on the *tidybayes, bayesplot*, and *patchwork* packages, as well as the *tidyverse* family of packages (Gabry *et al*., 2019; Kay, 2019; Pedersen, 2019; Wickham *et al*., 2019).

We used a multivariate generalized linear mixed model (GLMM) framework (Dingemanse & Dochtermann, 2013), which allowed us not only to draw inferences on dispersal and activity, but also on their correlations with each other and with (relative) shell mass, while handling missing shell mass data by in-model imputation (McElreath, 2020). A full write-up of the model is given in **Supplementary Material 2**. Briefly, we analysed the probability of dispersing and the probability of activity as individual-level binary variables (dispersed/did not disperse; active/not active) using Bernoulli GLMMs with a logit link. We tested whether dispersal or activity state depended on experienced population density, sex and body size (defined as ln(shell height); **Fig. 1**). The models also included random effects of date, experimental box, as well as a random effect of individual identity. Importantly, the latter was set to be identical (same values) in both the dispersal and activity models. In effect, this means it corresponds to a latent individual “behaviour” variable that would influence both dispersal and activity. This allowed us to estimate the effect of dispersal on activity (and vice-versa), while accounting for the fact both are estimated with uncertainty and the causal path from one to the other is unknown. We can then easily obtain the (logit) activity probability for, for instance, an average disperser by adding (on the logit scale) the average value of the latent behavioural variable for a disperser to the overall average logit(activity probability). The same principle applies to get the average dispersal probability of an active individual. Note that using instead dispersal status as a covariate in a univariate activity model leads to the exact same inferences (**Supplementary Material 3**); it however prevents us from doing the analyses described in the next few sentences. To be able to separate the effects of relative shell mass from those of body size, we used a third submodel. We fitted a linear mixed model in which ln(Shell mass) was a function of ln(Shell height), sex, and a random effect of individual identity. The use of repeated measures (see “Post-dispersal phenotyping”) of shell mass allowed us here to separate individual variation from residual error, and facilitated the estimation of the former. Finally, individual-level random effects were included in a variance-covariance matrix, allowing us to evaluate the correlation between “relative” shell mass (i.e. after accounting for body size and sex) and the latent behavioural variable. All continuous variables were centred and scaled to unit standard deviation (**Supplementary Material 4**), and the “sex” variable was converted to a centred dummy numerical variable (−0.5 = “Male”; +0.5 = “Female”)(Schielzeth, 2010).

We set weakly informative priors by following McElreath (2020)’s suggestions for continuous and proportion variables. We used Normal(μ = 0, σ = 1.5) priors for the dispersal and activity fixed effects intercepts, and Normal(0, 1) priors for all the other fixed effects. We set half-Normal(0, 1) priors for all standard deviations (random effects as well as residual variance of shell mass), and LKJ(η = 2) priors for the random effects correlation matrices.

We ran four chains for 25000 iterations, with the first 5000 of each chain used for warmup. We checked mixing graphically and confirmed chain convergence using the improved 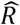 statistic (Vehtari *et al*., 2020). The chains were run longer than the default number of iterations (2000), to ensure effective sample size was satisfactory for all parameters (bulk- and tail-ESS sensu Vehtari *et al*., 2020 > 1000). All posterior summaries are given as: mean [95% Higher Posterior Density Intervals].

## Results

144 snails out of 371 (38.81%) dispersed across all test boxes. There was no detectable effect of box population density (**Fig. 2A**), sex or shell height on emigration probability (**Table 1**).

**Table 1.** Posterior means [and 95% credible intervals] for the parameters of the multivariate model of dispersal, activity and shell mass. Individual-level random effect covariance matrices are in **Table 2**. See **Supplementary Material 2** for the full write-up of the model, and **Supplementary Material 4**, for original means and SDs of transformed variables (*N* = 371 snails, except for Shell mass, *N* = 347 snails, each measured twice)

|  | Dispersal probability | Activity probability | Shell mass (ln-transformed then scaled) |
| --- | --- | --- | --- |
| *Fixed effects* |  |  |  |
| Intercept | -0.54 [-1.04 ; -0.26] | 0.74 [0.05 ; 1.49] | -0.01 [-0.09 ; 0.06] |
| Population density (scaled) | 0.04 [-0.38 ; 0.46] | -0.55 [-1.01 ; -0.10] | -- |
| Sex (effect of being female) | -0.16 [-0.79 ; 0.46] | 0.23 [-0.42 ; 0.86] | 0.18 [-0.02 ; 0.38] |
| Body size (shell height, ln-transformed then scaled) | 0.01 [-0.31 ; 0.33] | 0.16 [-0.17 ; 0.51] | 0.62 [0.52 ; 0.72] |
| *Random effects (see Table 2 for individual-level effects)* |  |  |  |
| Date-level variance | 0.13 [0.00 ; 0.59] | 0.56 [0.00 ; 1.75] | -- |
| Box-level variance | 0.52 [0.00 ; 1.33] | 0.38 [0.00 ; 1.04] | -- |
| Date-level dispersal-activity correlation | 0.13 [-0.69 ; 0.90] |  | -- |
| Box-level dispersal-activity correlation | 0.03 [-0.74 ; 0.73] |  | -- |
| *Residual variance* | -- | -- | $8.00 \times 10^{-4}$ [$6.85 \times 10^{-4}$ ; $9.23 \times 10^{-4}$] |

**Figure 2.**
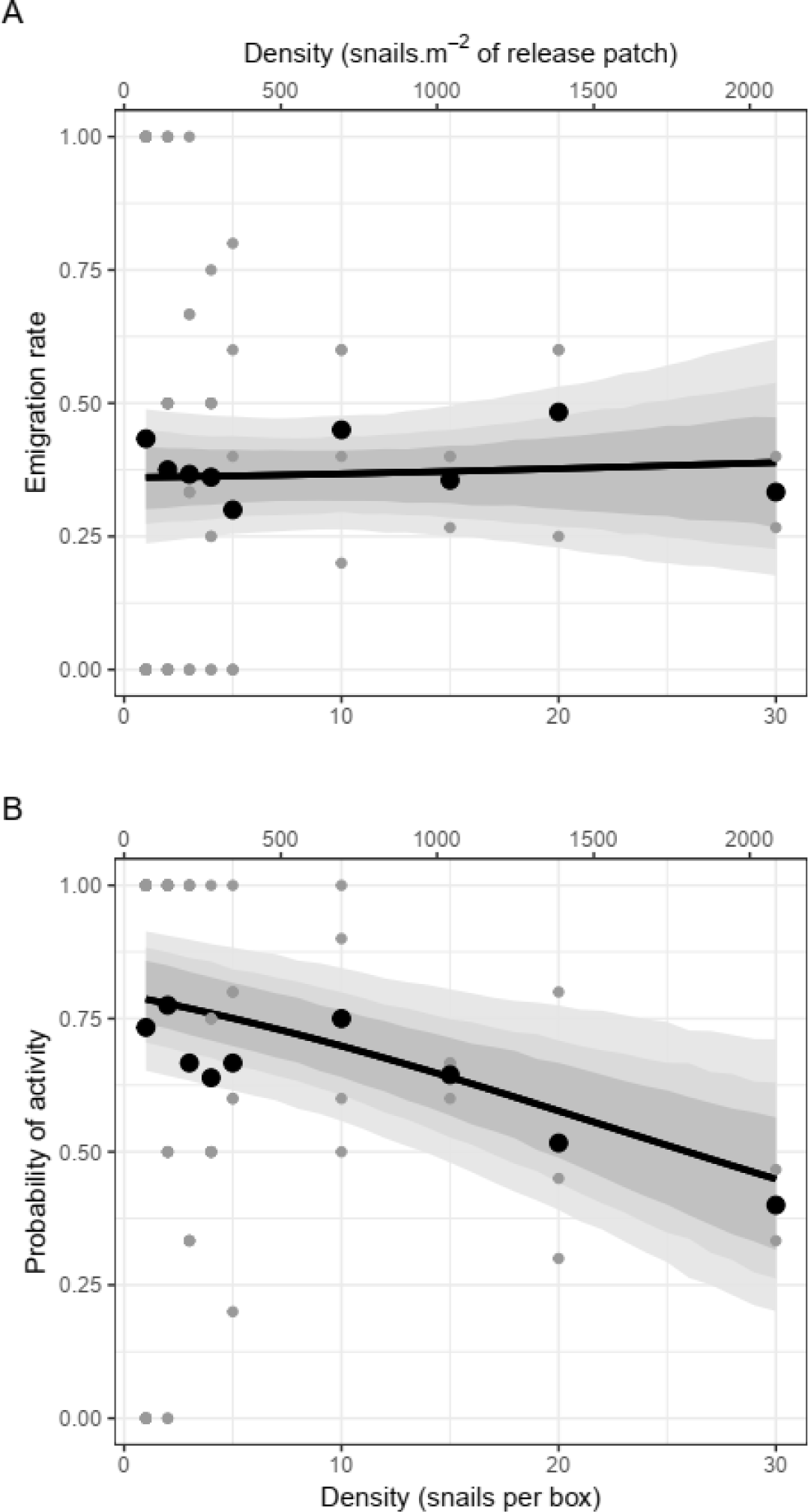
Effect of population density on *Pomatias elegans* dispersal rate (A) and probability of activity (B) (*N* = 371 snails). Grey points are observed proportions for each test box, black points averages for each density; black curves give posterior predicted means, grey bands 66, 84 and 95% credible intervals.

Snails exposed to higher population densities were less likely to be active afterwards (**Table 1, Fig. 2B**). There were no clear effects of sex or size on activity probability (**Table 1**). The latent behavioural variable was positively associated with both emigration and activity (**Supplementary Material 5**): accordingly, dispersers were 1.23 [1.02; 1.47] times more likely to be active on average, and conversely active individuals were 1.54 [1.04; 2.16] times more likely to have emigrated (**Fig. 3**). This activity-emigration correlation was confirmed by analysing activity in a univariate model with dispersal status as a covariate (**Supplementary Material 3**). After accounting for shell length, sex and measurement error, individual shell mass was negatively correlated with the behavioural latent variable: dispersers/active snails tended to have lighter shells, given their size and sex, than residents/inactive individuals (**Fig. 4, Table 2**).

**Table 2.** Individual-level variance-covariance matrix between the latent behavioural variable (positively correlated with dispersal and activity, see **Fig. 3** and **Supplementary Material 3**) and shell mass. Posterior means [95% credible interval] are depicted; random effects variances are given on the diagonal, correlation/covariance on the off diagonal

|  | Behaviour latent variable | Shell mass |
| --- | --- | --- |
| Behaviour latent variable | 0.90 [0.08 ; 1.81] |  |
| Shell mass | -0.39 [-0.67 ; -0.12] / -0.25 [-0.41 ; -0.09] | 0.54 [0.46 ; 0.63] |

**Figure 3.**
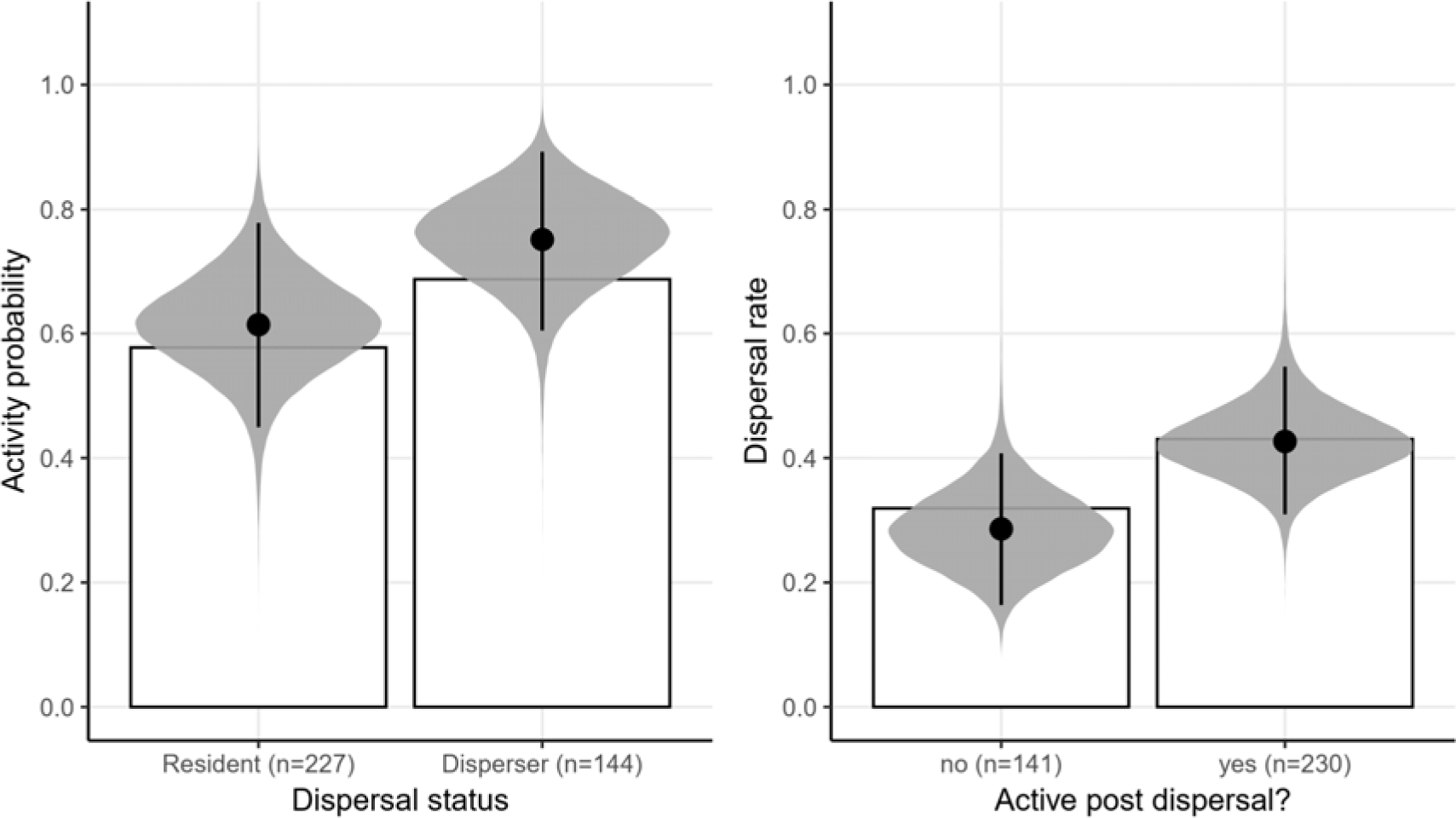
Relationship between dispersal status and activity post-dispersal. White bars correspond to observed frequencies, grey “eyes” to posterior predictions based on model intercepts and individual-level random effects, i.e. averaging out the effects of sex, size and population density (black dots: posterior means; segments: 95% credible intervals).

**Figure 4.**
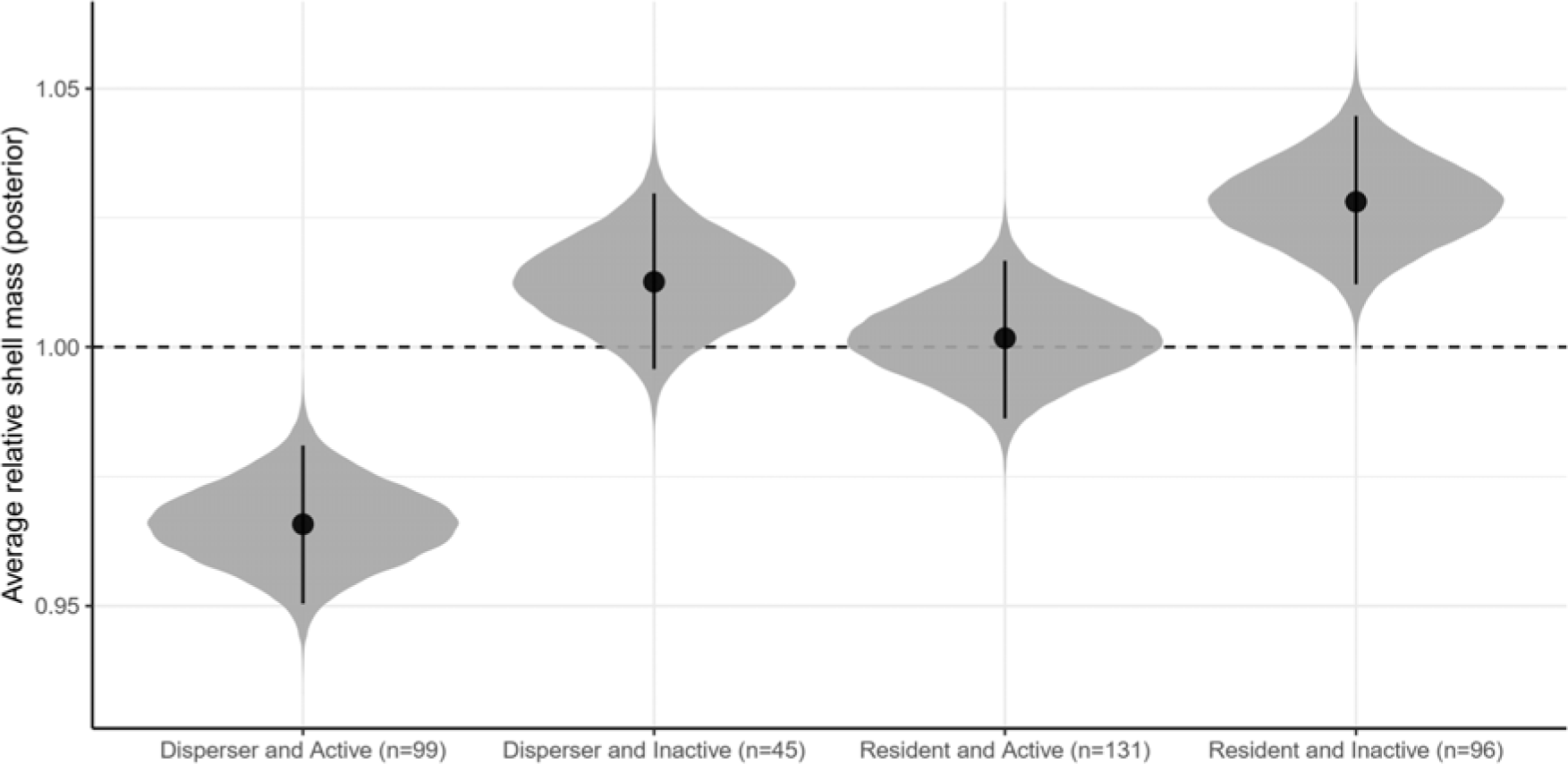
Posterior predicted relationship between dispersal status, activity post-dispersal, and mean predicted shell mass, that is relative to the shell mass predicted for a snail of same shell height and sex (black dots: posterior means; segments: 95% credible intervals). Posterior pairwise mean differences are presented in **Supplementary Material 6**.

## Discussion

We studied the relationship between *Pomatias elegans* dispersal and population density over most of the known range of natural densities (Pfenninger, 2002; Albano *et al*., 2015), yet found no evidence of density-dependent emigration. There was however a negative relationship between density and activity. Furthermore, we found that emigration, activity and (relative) shell mass co-vary in this species, once density variation is accounted for.

We additionally found no evidence of sex bias in dispersal probability or activity. While sex-biased dispersal is abundantly documented in non-molluscan groups (Bowler & Benton, 2005; Trochet *et al*., 2016), there are numerous species with no bias. Broadly, sex-dependent dispersal is predicted to arise when there are sex-differences in spatio-temporal fitness variation due to e.g. sex differences in resource use and/or in local mate competition (Hovestadt *et al*., 2014; Trochet *et al*., 2016; Li & Kokko, 2019). Our results, in light of dispersal theory, imply that male and female *P. elegans* use resources similarly and that they do not present polygyny or any mating system leading to predictable sex differences in spatio-temporal fitness variation. While sex ratio is, overall, balanced in this species (Boycott, 1916), the other predictions remain to be tested by the collection of further natural history data. We should additionally note that our experimental setup (one measure of dispersal, after 4 days, and groups that were sex-balanced on average) may not have allowed us to detect finer sex differences in dispersal, i.e. in emigration timing (Li & Kokko, 2019), or conditional strategies dependent on locally experienced sex-ratio (Hovestadt *et al*., 2014).

The absence of density-dependent emigration was more surprising, as many other land mollusc species have been shown to increase emigration and/or dispersal distances in response to population density (Cain & Currey, 1968; Oosterhoff, 1977; Hamilton & Wellington, 1981; Livshits, 1985; Baur, 1993; Shachak, Leeper & Degen, 2002; Dahirel *et al*., 2016). In addition, our experimental design included a (supposedly) costly matrix, which should have made the detection of a positive emigration-density link easier, if it exists (as the dispersal-density reaction norm is predicted to become steeper as costs of dispersal increase; Travis *et al*., 1999). The costs of competition, which can trigger density-dependent dispersal, are well-documented in land molluscs (Oosterhoff, 1977; Hamilton & Wellington, 1981; Dan & Bailey, 1982). However, density-independent emigration also exists in some mollusc species, in at least some contexts (e.g. Hamilton & Wellington, 1981; Baur, 1993). The traditionally expected positive relationship between density and dispersal can be dampened, or even reversed, if for instance species/individuals benefit from group-living (Serrano *et al*., 2003; Bowler & Benton, 2005; Cote & Clobert, 2007). In terrestrial gastropods, individuals may aggregate actively in shelters (and get benefits from this aggregation that go beyond the shelter itself, as the presence of conspecifics may create a favourable microclimate), sometimes following trails to do so (Cook, 2001). However, it is unknown whether *Pomatias elegans* follows trails, and this aggregation behaviour is anyway common to many land molluscs, including species we know show positive density-dependent emigration (Cook, 2001), so it is unlikely to explain alone the density-independence seen in the present study. *P. elegans*, contrary to most land molluscs has separate sexes, and differences in reproductive systems may drive the evolution of different dispersal strategies (Eppley & Jesson, 2008). Indeed, all things equal, hermaphrodites are *a priori* more likely than species with separate sexes to find a mate in small populations (Tomlinson, 1966). This means the density threshold at which the costs of competition outweigh the benefits of grouping (Kim, Torres & Drummond, 2009; Fronhofer, Kropf & Altermatt, 2015) and the costs of dispersal (Bonte *et al*., 2012) is likely lower for hermaphrodites. We hypothesize that context being equal (including e.g. movement costs, or group shelter benefits), this may make hermaphrodites more likely than gonochoric species to exhibit positive density-dependent dispersal over a range of natural densities. In any case, our study shows that further comparisons between hermaphroditic and gonochoric species sharing ecological and life-history characteristics are crucially needed to clarify the role of reproductive system in shaping context-dependent dispersal.

Alternatively (but not exclusively), the absence of density response in dispersal may reflect the fact *Pomatias* snails possess other mechanisms to respond to unfavourable densities. Indeed, by contrast with dispersal, activity propensity did decrease with increasing population density (**Fig. 2B**). Like many other shelled land gastropods, *P. elegans* can reduce its activity and even enter dormancy when context is unfavourable (Cook, 2001; Falkner *et al*., 2001). Activity reduction in response to increased competition has been observed in several species (Oosterhoff, 1977; Cameron & Carter, 1979; Dan & Bailey, 1982). This is generally thought to result from interference (mediated through mucus and/or faeces) by the most competitive individuals, which remain active. However, when the experimental/observational setup also allowed for dispersal, species that reduced activity in response to density (e.g. *Cepaea nemoralis, Cornu aspersum, Xerocrassa seetzenii*) did however also exhibit density-dependent dispersal (Oosterhoff, 1977; Shachak *et al*., 2002; Dahirel *et al*., 2016). Given the very limited dispersal capacities of *P. elegans* (Pfenninger, 2002) even compared to other snails (Kramarenko, 2014), the least competitive *Pomatias* snails may well be more successful reducing activity to wait out more favourable conditions, than by attempting to find better by moving away. Indeed, in snails, starvation, a possible result of inactivity, may trigger entry into actual dormancy (Wells, 1944), which can be seen as a form of “dispersal in time” (Buoro & Carlson, 2014). The fact there was no effect of body size on activity suggests the loss of activity is not simply an inhibition of activity by competitive individuals, leading to competitive exclusion. Indeed, in that case we would have expected larger individuals to stay active at high densities while smaller less competitive snails are inhibited (Dan & Bailey, 1982). Given this, and while we need more information to conclude that strongly reduced activity in short-term tests is correlated with long-term dormancy propensity, it is possible that *Pomatias* snails use dormancy as a general “dispersal in time” mechanism, rather than moving away in response to high densities. Note however that we only studied emigration, the first step of dispersal, and density-dependent responses may also happen during transience (Shachak *et al*., 2002; Poethke, Gros & Hovestadt, 2011; Dahirel *et al*., 2016). In addition, we set within-patch conditions to be standard and favourable; food availability may interact with density and shape dispersal (e.g. Endriss *et al*., 2019).

All three phenotypic traits we studied were linked. First, density being held equal, dispersers were more likely to be active post-dispersal test, and vice versa (**Fig. 3, Supplementary Material 3**). These results line up with predictions on dispersal syndromes and the pace-of-life hypothesis (Cote *et al*., 2010; Réale *et al*., 2010), which predicts among other things a metabolically-underpinned link between activity and risk taking behaviour (such as dispersal). While, again, more extended assessments of activity are needed to confirm that low activity is correlated to true dormancy, the observed syndrome can also be interpreted as a trade-off between dispersal in space and in time (Buoro & Carlson, 2014), in which individuals that do not disperse are more likely to be dormant at any point in time, even after accounting for density effects. Second, we found that more mobile snails (higher activity and/or dispersal) had on average lighter shells (**Fig. 4, Table 2**), compared to inactive/resident snails. This result goes against our predictions based on the phenotypic compensation hypothesis, which states that risk-taking individuals should have better morphological defences against predators (Hulthén *et al*., 2014; Ahlgren *et al*., 2015; Kuo, Irschick & Lailvaux, 2015). However, phenotypic compensation is not a hard rule (De Winter *et al*., 2016), and while thicker/heavier shells are harder to break (see e.g. Rosin *et al*., 2013), variation in shell characteristics depends on many abiotic and biotic parameters (Goodfriend, 1986). Indeed, thicker shells may also better conserve humidity (Goodfriend, 1986), which may be adaptive for snails more prone to be inactive for extended periods of time. Heavier shells are likely to increase the costs of movement (Herreid & Full, 1986); given the already high costs of movement in land molluscs (Denny, 1980), this may explain why mobile individuals have lighter shells, despite the potentially increased predation risk. Experimental manipulations of the costs and benefits of heavier shells (by e.g. testing snails under varying humidity conditions, and/or by adding weights to shells), may help disentangle the underpinnings of this dispersal/shell mass syndrome.

Although *P. elegans* is overall much less abundant than other macroinvertebrate decomposers at the landscape scale (David, 1999), it can locally reach very high densities (Pfenninger, 2002). Our study shows that increased densities do not trigger increased emigration from crowded into empty habitats in this species. This helps explain the maintenance of this strong spatial heterogeneity and high local densities, as dispersal cannot play its “density-equalizing” role as well. Given decomposition by *P. elegans* can have marked effects on litter characteristics (Coulis *et al*., 2009; De Oliveira *et al*., 2010), and facilitate decomposition by other co-occurring macroinvertebrates (De Oliveira *et al*., 2010), this absence of a common context-dependency in dispersal may have broader implications for local heterogeneity in soil ecosystem functioning and soil communities.

## Acknowledgements

We thank Valérie Briand for her help in obtaining some of the cited scientific articles, as well as the editor and two anonymous reviewers, for helpful comments on a previous version of this manuscript.

## Data availability

Data and analysis script are archived on GitHub (https://github.com/mdahirel/pomatias-dispersal-2017) and Zenodo (doi: 10.5281/zenodo.3691977) (version v1.2).

## References

Ahlgren, J., Chapman, B.B., Nilsson, P.A. & Brönmark, C. (2015). Individual boldness is linked to protective shell shape in aquatic snails. Biol. Lett.11, 20150029.

Albano, P.G., Strazzari, G., D’Occhio, P. & Succetti, F. (2015). Field estimates of detectability and site occupancy show that northern Italy forest molluscs are spatially rare and poorly detectable. Ital. J. Zool.82, 592–608.

Baines, C.B., Ferzoco, I.M.C. & McCauley, S.J. (2019). Phenotype-by-environment interactions influence dispersal. J. Anim. Ecol.88, 1263–1274.

Barker, G.M. (2001). Gastropods on land: phylogeny, diversity and adaptive morphology. In The biology of terrestrial molluscs: 1–146. Barker, G.M. (Ed.). Wallingford, UK: CABI.

Baur, A. (1993). Effects of food availability and intra-and interspecific interaction on the dispersal tendency in the land snail Chondrina clienta. J. Zool.230, 87–100.

Bonte, D. & Dahirel, M. (2017). Dispersal: a central and independent trait in life history. Oikos 126, 472–479.

Bonte, D., Van Dyck, H., Bullock, J.M., Coulon, A., Delgado, M., Gibbs, M., Lehouck, V., Matthysen, E., Mustin, K., Saastamoinen, M., Schtickzelle, N., Stevens, V.M., Vandewoestijne, S., Baguette, M., Barton, K., Benton, T.G., Chaput-Bardy, A., Clobert, J., Dytham, C., Hovestadt, T., Meier, C.M., Palmer, S.C.F., Turlure, C. & Travis, J.M.J. (2012). Costs of dispersal. Biol. Rev.87, 290–312.

Bowler, D.E. & Benton, T.G. (2005). Causes and consequences of animal dispersal strategies: relating individual behaviour to spatial dynamics. Biol. Rev.80, 205–225.

Boycott, A.E. (1916). On sexual characteristics in the shell and radula of Pomatias elegans (Müller). Proc. Malacol. Soc. Lond. v.12-13 (1916-1919), 127–132.

Buoro, M. & Carlson, S.M. (2014). Life-history syndromes: Integrating dispersal through space and time. Ecol. Lett.17, 756–767.

Bürkner, P.-C. (2017). brms: an R package for Bayesian multilevel models using Stan. J. Stat. Softw.80, 1–28.

Cain, A.J. & Currey, J.D. (1968). Ecogenetics of a population of Cepaea nemoralis (L.) subject to strong area effects. Phil. Trans. R. Soc. B 253, 447–482.

Cameron, R.A.D. & Carter, M.A. (1979). Intra- and interspecific effects of population density on growth and activity in some helicid land snails (Gastropoda: Pulmonata). J. Anim. Ecol.48, 237–246.

Carpenter, B., Gelman, A., Hoffman, M.D., Lee, D., Goodrich, B., Betancourt, M., Brubaker, M., Guo, J., Li, P. & Riddell, A. (2017). Stan: a probabilistic programming language. J. Stat. Softw.76, 1–32.

Clobert, J., Baguette, M., Benton, T.G. & Bullock, J.M. (Eds.). (2012). Dispersal ecology and evolution. Oxford, UK: Oxford University Press.

Cook, A. (2001). Behavioural ecology: on doing the right thing, in the right place at the right time. In The biology of terrestrial molluscs: 447–488. Barker, G.M. (Ed.). Wallingford, UK: CABI.

Cote, J. & Clobert, J. (2007). Social personalities influence natal dispersal in a lizard. Proc. Royal Soc. B 274, 383–390.

Cote, J., Clobert, J., Brodin, T., Fogarty, S. & Sih, A. (2010). Personality-dependent dispersal: characterization, ontogeny and consequences for spatially structured populations. Phil. Trans. R. Soc. B 365, 4065–4076.

Coulis, M., Hättenschwiler, S., Rapior, S. & Coq, S. (2009). The fate of condensed tannins during litter consumption by soil animals. Soil Biol. Biochem.41, 2573–2578.

Dahirel, M., Vardakis, M., Ansart, A. & Madec, L. (2016). Density-dependence across dispersal stages in a hermaphrodite land snail: insights from discrete choice models. Oecologia 181, 1117–1128.

Dan, N. & Bailey, S.E.R. (1982). Growth, mortality, and feeding rates of the snail Helix aspersa at different population densities in the laboratory, and the depression of activity of helicid snails by other individuals, or their mucus. J. Mollus. Stud.48, 257–265.

David, J.-F. (1999). Abundance, biomass and functional structure of the saprophagous macrofauna in the litter and soil of Mediterranean oak forests. Pedobiologia 43, 319–327.

De Meester, N. & Bonte, D. (2010). Information use and density-dependent emigration in an agrobiont spider. Behav. Ecol.21, 992–998.

De Oliveira, T., Hättenschwiler, S. & Handa, I.T. (2010). Snail and millipede complementarity in decomposing Mediterranean forest leaf litter mixtures. Funct. Ecol.24, 937–946.

De Winter, G., Ramalho Martins, H., Trovo, R.A. & Chapman, B.B. (2016). Knights in shining armour are not necessarily bold: defensive morphology correlates negatively with boldness, but positively with activity, in wild threespine stickleback, Gasterosteus aculeatus. Evol. Ecol. Res.17, 1–12.

Denny, M. (1980). Locomotion: the cost of gastropod crawling. Science 208, 1288–1290.

Dingemanse, N.J. & Dochtermann, N.A. (2013). Quantifying individual variation in behaviour: mixed-effect modelling approaches. J. Anim. Ecol.82, 39–54.

Endriss, S.B., Vahsen, M.L., Bitume, E.V., Monroe, J.G., Turner, K.G., Norton, A.P. & Hufbauer, R.A. (2019). The importance of growing up: juvenile environment influences dispersal of individuals and their neighbours. Ecol. Lett.22, 45–55.

Eppley, S.M. & Jesson, L.K. (2008). Moving to mate: the evolution of separate and combined sexes in multicellular organisms. J. Evol. Biol.21, 727–736.

Falkner, G., Obrdlik, P., Castella, E. & Speight, M.C.D. (2001). Shelled Gastropoda of western Europe. München, Germany: Friedrich Held Gesellschaft.

Fronhofer, E.A., Kropf, T. & Altermatt, F. (2015). Density-dependent movement and the consequences of the Allee effect in the model organism Tetrahymena. J. Anim. Ecol.84, 712–722.

Fronhofer, E.A., Legrand, D., Altermatt, F., Ansart, A., Blanchet, S., Bonte, D., Chaine, A., Dahirel, M., Laender, F.D., Raedt, J.D., Gesu, L. di, Jacob, S., Kaltz, O., Laurent, E., Little, C.J., Madec, L., Manzi, F., Masier, S., Pellerin, F., Pennekamp, F., Schtickzelle, N., Therry, L., Vong, A., Winandy, L. & Cote, J. (2018). Bottom-up and top-down control of dispersal across major organismal groups. Nature Ecol. Evol.2, 1859–1863.

Gabry, J., Simpson, D., Vehtari, A., Betancourt, M. & Gelman, A. (2019). Visualization in Bayesian workflow. J. Royal Stat. Soc. A 182, 389–402.

Goodfriend, G.A. (1986). Variation in land-snail shell form and size and its causes: a review. Syst. Zool.35, 204–223.

Hamilton, P.A. & Wellington, W.G. (1981). The effects of food and density of the movement of Arion ater and Ariolimax columbianus (Pulmonata: Stylommatophora) between habitats. Res. Popul. Ecol.23, 299–308.

Harman, R.R., Goddard, J., Shivaji, R. & Cronin, J.T. (2020). Frequency of occurrence and population-dynamic consequences of different forms of density-dependent emigration. Am. Nat.195, 851–867.

Herreid, C.F. & Full, R.J. (1986). Energetics of hermit crabs during locomotion: the cost of carrying a shell. J. Exp. Biol.120, 297–308.

Hovestadt, T., Mitesser, O. & Poethke, H.-J. (2014). Gender-specific emigration decisions sensitive to local male and female density. Am. Nat.184, 38–51.

Hulthén, K., Chapman, B.B., Nilsson, P.A., Hollander, J. & Brönmark, C. (2014). Express yourself: bold individuals induce enhanced morphological defences. Proc. R. Soc. B 281, 20132703.

Jacob, S., Chaine, A.S., Huet, M., Clobert, J. & Legrand, D. (2019). Variability in dispersal syndromes is a key driver of metapopulation dynamics in experimental microcosms. Am. Nat.194, 613–626.

Kay, M. (2019). tidybayes: tidy data and geoms for Bayesian models.

Kerney, M.-P. & Cameron, R.-A.-D. (1999). Guide des escargots et limaces d’Europe. Lonay (Suisse): Delachaux et Niestlé.

Kilian, E.F. (1951). Untersuchungen zur Biologie von Pomatias elegans (Müller) und ihrer ‘Konkrementdrüse.’ Arch. Molluskenkd.80, 1–16.

Kim, S.-Y., Torres, R. & Drummond, H. (2009). Simultaneous positive and negative density-dependent dispersal in a colonial bird species. Ecology 90, 230–239.

Kramarenko, S. (2014). Active and passive dispersal of terrestrial mollusks: a review. Ruthenica 24, 1–14.

Kuo, C.-Y., Irschick, D.J. & Lailvaux, S.P. (2015). Trait compensation between boldness and the propensity for tail autotomy under different food availabilities in similarly aged brown anole lizards. Funct. Ecol.29, 385–392.

Li, X.-Y. & Kokko, H. (2019). Intersexual resource competition and the evolution of sex-biased dispersal. Front. Ecol. Evol.7, 111.

Livshits, G.M. (1985). Ecology of the terrestrial snail Brephulopsis bidens (Pulmonata: Enidae): mortality, burrowing and migratory activity. Malacologia 26, 213–223.

Matthysen, E. (2005). Density-dependent dispersal in birds and mammals.Ecography 28, 403–416.

Matthysen, E. (2012). Multicausality of dispersal: a review. In Dispersal ecology and evolution: 3–18.

Clobert, J., Baguette, M., Benton, T.G. & Bullock, J.M. (Eds.). Oxford, UK: Oxford University Press.

McElreath, R. (2020). Statistical rethinking: a Bayesian course with examples in R and Stan. 2nd edition. Boca Raton, USA: Chapman and Hall/CRC.

Oosterhoff, L.M. (1977). Variation in growth rate as an ecological factor in the landsnail Cepaea nemoralis (L.). Neth. J. Zool.27, 1–132.

Pedersen, T.L. (2019). patchwork: the composer of plots.

Pfenninger, M. (2002). Relationship between microspatial population genetic structure and habitat heterogeneity in Pomatias elegans (O.F. Müller 1774) (Caenogastropoda, Pomatiasidae). Biol. J. Linn. Soc.76, 565–575.

Poethke, H.J., Gros, A. & Hovestadt, T. (2011). The ability of individuals to assess population density influences the evolution of emigration propensity and dispersal distance. Theor. Biol.282, 93–99.

R Core Team. (2020). R: a language and environment for statistical computing. Vienna, Austria: R Foundation for Statistical Computing.

Réale, D., Garant, D., Humphries, M.M., Bergeron, P., Careau, V. & Montiglio, P.-O. (2010). Personality and the emergence of the pace-of-life syndrome concept at the population level. Phil. Trans. R. Soc. B 365, 4051–4063.

Rodrigues, A.M.M. & Johnstone, R.A. (2014). Evolution of positive and negative density-dependent dispersal. Proc. R. Soc. B 281, 20141226.

Ronce, O. (2007). How does it feel to be like a rolling stone? Ten questions about dispersal evolution. Annu. Rev. Ecol. Evol. Syst.38, 231–253.

Rosin, Z.M., Kobak, J., Lesicki, A. & Tryjanowski, P. (2013). Differential shell strength of Cepaea nemoralis colour morphs—implications for their anti-predator defence. Naturwissenschaften 100, 843–851.

Schielzeth, H. (2010). Simple means to improve the interpretability of regression coefficients. Methods Ecol. Evol.1, 103–113.

Serrano, D., Tella, J.L., Donázar, J.A. & Pomarol, M. (2003). Social and individual features affecting natal dispersal in the colonial lesser kestrel. Ecology 84, 3044–3054.

Shachak, M., Leeper, A. & Degen, A. (2002). Effect of population density on water influx and distribution in the desert snail Trochoidea seetzenii. Ecoscience 9, 287–292.

Stan Development Team. (2018). RStan: the R interface to Stan.Tomlinson, J. (1966). The advantages of hermaphroditism and parthenogenesis. J. Theor. Biol.11, 54–58.

Travis, J.M.J., Murrell, D.J. & Dytham, C. (1999). The evolution of density–dependent dispersal. Proc. R. Soc. B 266, 1837–1842.

Trochet, A., Courtois, E.A., Stevens, V.M., Baguette, M., Chaine, A., Schmeller, D.S., Clobert, J. & Wiens, J.J. (2016). Evolution of sex-biased dispersal. Q. Rev. Biol.91, 297–320.

Van Dyck, H. & Baguette, M. (2005). Dispersal behaviour in fragmented landscapes: routine or special movements? Basic Appl. Ecol.6, 535–545.

Vehtari, A., Gelman, A., Simpson, D., Carpenter, B. & Bürkner, P.-C. (2020). Rank-normalization, folding, and localization: an improved \widehat{R} for assessing convergence of MCMC. Bayesian Anal.

Wells, G.P. (1944). The water relations of snails and slugs: III. Factors determining activity in Helix pomatia L. J. Exp. Biol.20, 79–87.

Welter-Schultes, F. (2012). European non-marine molluscs, a guide for species identification. Göttingen: Planet Poster Editions.

Wickham, H., Averick, M., Bryan, J., Chang, W., McGowan, L., François, R., Grolemund, G., Hayes, A., Henry, L., Hester, J., Kuhn, M., Pedersen, T., Miller, E., Bache, S., Müller, K., Ooms, J., Robinson, D., Seidel, D., Spinu, V., Takahashi, K., Vaughan, D., Wilke, C., Woo, K. & Yutani, H. (2019). Welcome to the Tidyverse. J. Open Source Softw.4, 1686.

