## Supplementary Material for "Increased population density depresses activity but does not influence emigration in the snail *Pomatias elegans*"

**Supplementary Material 1: sexing snails, and confirming the lack of sex bias in the final dataset**

We aimed to have balanced sex ratios across densities (in line with natural populations ; Boycott, 1916). However, reproductive organs are not externally visible in *Pomatias elegans*, and body size is not an accurate enough correlate, meaning a classification in males and females can only be done by dissection (Creek, 1951). We nonetheless used the fact that females are on average slightly bigger and wider than males (Boycott, 1916; Jordaens, Platts & Backeljau, 2001) to putatively sex snails before the experiment, and then assigned (roughly) equal numbers of putative males and females to densities and boxes. To predict sex, we first measured the shell height, shell width and aperture height (**Fig. 1** in main text) to the nearest 0.1 mm in a set of 26 individuals not used in dispersal experiments, and sexed them by dissection after freezing at -20°C. We used these individuals to fit an initial Linear Discriminant Analysis (LDA) predicting sex as a function of shell characteristics, and used this LDA to predict the sex of our focal snails. Experimental individuals were then added to the LDA dataset once dissected after each test session, in order to improve its predictive ability. Overall, 96 individuals out of 371 (25.88%) were misclassified prior to experiments (49 out of 185 males, based on reproductive organs, were first classified as females, 47 females out of 186 first classified as males).

To evaluate whether our snail sample ($N$ = 371 in 87 boxes) was biased with respect to sex, we ran two binomial generalized linear models, with $\mathrm{Normal}(\mu = 0, \sigma = 1.5)$ priors for all coefficients (as they correspond to logit(proportions); McElreath, 2020). The first one was an intercept-only model, to determine whether there was an overall sex bias. The second model included population density as a categorical variable, to determine whether specific densities showed sex biases. In both cases, we found no clear evidence of sex bias, see **Table S1** next page.

Supplementary Material for: "Increased population density depresses activity but does not influence emigration in the snail *Pomatias elegans*", by Maxime Dahirel, Loïc Menut, Armelle Ansart

**Table S1: predicted proportion of females in the entire dataset and at each tested density**

| Density (snails per box) | Number of replicates (boxes) | Predicted proportion of females; posterior mean [95% credible interval] |
|---|---|---|
| All (intercept only model) | 87 | 0.50 [0.45 ; 0.55] |
| 1 | 30 | 0.50 [0.33 ; 0.68] |
| 2 | 20 | 0.60 [0.45; 0.74] |
| 3 | 10 | 0.53 [0.37; 0.69] |
| 4 | 9 | 0.42 [0.27; 0.58] |
| 5 | 6 | 0.60 [0.43; 0.76] |
| 10 | 4 | 0.45 [0.31; 0.61] |
| 15 | 3 | 0.53 [0.39; 0.66] |
| 20 | 3 | 0.55 [0.43; 0.66] |
| 30 | 2 | 0.39 [0.27; 0.50] |

**Supplementary Material 2: detailed description of the main multivariate model**

The model analysed in the main text can be written as following:

$$\text{Dispersal}_{i,j,t} \sim \text{Bernoulli}(p_{i,j,t})$$

$$\text{Activity}_{i,j,t} \sim \text{Bernoulli}(q_{i,j,t})$$

$$\text{Z}(\ln(\text{Shell Mass}))_{i,j,k} \sim \text{Normal}(\mu_{i,j}, \sigma_r)$$

$$logit(p_{i,j,t}) = \beta_{0[\text{dispersal}]} + \beta_{1[\text{dispersal}]} \times \text{Z}(\text{Density})_i + \beta_{2[\text{dispersal}]} \times \text{Z}(\ln(\text{Shell Height}))_{i,j} + \beta_{3[\text{dispersal}]} \times \text{Sex}_{i,j} + \alpha_{[\text{dispersal}]i} + \gamma_{[\text{dispersal}]t} + \eta_{[\text{behaviour}]j}$$

$$logit(q_{i,j,t}) = \beta_{0[\text{activity}]} + \beta_{1[\text{activity}]} \times \text{Z}(\text{Density})_i + \beta_{2[\text{activity}]} \times \text{Z}(\ln(\text{Shell Height}))_{i,j} + \beta_{3[\text{activity}]} \times \text{Sex}_{i,j} + \alpha_{[\text{activity}]i} + \gamma_{[\text{activity}]t} + \eta_{[\text{behaviour}]j}$$

$$\mu_{i,j} = \beta_{0[\text{shell mass}]} + \beta_{1[\text{shell mass}]} \times \text{Z}(\ln(\text{Shell Height}))_{i,j} + \beta_{2[\text{shell mass}]} \times \text{Sex}_{i,j} + \eta_{[\text{shell mass}]j}$$

, in which $p_{i,j,t}$ and $q_{i,j,t}$ are dispersal and activity probabilities of individual *j* in box *i,* whose test *started* at date *t* (so activity and dispersal from the individual are coded with the same date); Z() is the function transforming a vector of values to their Z-scores; i.e. mean-centring and scaling to unit

1SD; the $_{\text{[dispersal]}}$, $_{\text{[activity]}}$ and $_{\text{[shell mass]}}$ subscripts denote the response variable a parameter corresponds to ($_{\text{[behaviour]}}$ indicates a parameter is common to both dispersal and activity); $\mathrm{Z}(\ln(\text{Shell Mass}))_{i,j,k}$ is the (ln-transformed and scaled) observed shell mass of individual *j* in box *i* during measurement *k,* $\mu_{i,j}$ the true mass (same scale) of the corresponding individual, $\sigma_r$ the measurement error; $\beta_h$ are the fixed effect coefficients, $\alpha_i$, $\gamma_t$ and $\eta_j$ the box- , date- , and individual-specific average deviations from fixed effect/ population-level estimates.

The random effects $\alpha$, $\gamma$ and $\eta$ are distributed according to the following:

$$\begin{bmatrix} \alpha_{\text{[dispersal]}i} \\ \alpha_{\text{[activity]}i} \end{bmatrix} \sim MVNormal(\begin{bmatrix} 0 \\ 0 \end{bmatrix}, \mathbf{\Omega_{box}})$$

$$\begin{bmatrix} \gamma_{\text{[dispersal]}t} \\ \gamma_{\text{[activity]}t} \end{bmatrix} \sim MVNormal(\begin{bmatrix} 0 \\ 0 \end{bmatrix}, \mathbf{\Omega_{date}})$$

$$\begin{bmatrix} \eta_{\text{[behaviour]}j} \\ \eta_{\text{[shell mass]}j} \end{bmatrix} \sim MVNormal(\begin{bmatrix} 0 \\ 0 \end{bmatrix}, \mathbf{\Omega_{ind}})$$

$$\mathbf{\Omega_{box}} = \begin{bmatrix} \sigma_{\alpha_{\text{[dispersal]}}} & 0 \\ 0 & \sigma_{\alpha_{\text{[activity]}}} \end{bmatrix} \mathbf{R_{box}} \begin{bmatrix} \sigma_{\alpha_{\text{[dispersal]}}} & 0 \\ 0 & \sigma_{\alpha_{\text{[activity]}}} \end{bmatrix}$$

$$\mathbf{\Omega_{date}} = \begin{bmatrix} \sigma_{\gamma_{\text{[dispersal]}}} & 0 \\ 0 & \sigma_{\gamma_{\text{[activity]}}} \end{bmatrix} \mathbf{R_{date}} \begin{bmatrix} \sigma_{\gamma_{\text{[dispersal]}}} & 0 \\ 0 & \sigma_{\gamma_{\text{[activity]}}} \end{bmatrix}$$

$$\mathbf{\Omega_{ind}} = \begin{bmatrix} \sigma_{\eta_{\text{[behaviour]}}} & 0 \\ 0 & \sigma_{\eta_{\text{[shell mass]}}} \end{bmatrix} \mathbf{R_{ind}} \begin{bmatrix} \sigma_{\eta_{\text{[behaviour]}}} & 0 \\ 0 & \sigma_{\eta_{\text{[shell mass]}}} \end{bmatrix}$$

, where $\mathbf{\Omega_{box}}$ , $\mathbf{\Omega_{date}}$ and $\mathbf{\Omega_{ind}}$ are the variance-covariance matrices for the box- and individual-level random effects, with $\mathbf{R_{box}}$ , $\mathbf{R_{date}}$ and $\mathbf{R_{ind}}$ the corresponding correlation matrices, and $\sigma$ the random effect standard deviations.

The associated prior distributions follow McElreath (2020) and were $\text{Normal}(\mu = 0, \sigma = 1.5)$ for the intercepts $\beta_0$ for dispersal and activity; $\text{Normal}(0,1)$ for all other fixed effects $\beta_h$; $\text{half} - \text{Normal}(0,1)$ for all standard deviations noted $\sigma$, including the residual standard deviation $\sigma_r$, and a LKJ prior $\text{LKJcorr}(\eta = 2)$ for the correlation matrices $\mathbf{R}$.

**Supplementary Material 3: using dispersal as covariate in the activity model**

To evaluate the link between dispersal and activity behaviour, one can alternatively use the following univariate model:

$$\text{Activity}_{i,j,t} \sim \text{Bernoulli}(q_{i,j,t})$$

$$logit(q_{i,j,t}) = \beta_{0[\text{activity}]} + \beta_{1[\text{activity}]} \times \text{Z(Density)}_{i} + \beta_{2[\text{activity}]} \times \text{Z(ln(Shell Height))}_{i,j} + \beta_{3[\text{activity}]} \times \text{Sex}_{i,j} + \beta_{4[\text{activity}]} \times \text{Dispersal}_{i,j} + \alpha_{[\text{activity}]i} + \gamma_{[\text{activity}]t}$$

(with Dispersal entered as a centered dummy variable, similarly to sex). Running this univariate model, with the same priors as in **Supplementary Material 2** (when relevant) leads to an estimate of $\beta_{4[\text{activity}]}$ = 0.55 [0.04 ; 0.99], confirming that, as shown in the main text, dispersers are indeed more likely to be active post-dispersal. The converse model, with emigration as the response and activity as the covariate, confirms the association between emigration and activity ($\beta_{4[\text{dispersal}]}$ = 0.56 [0.08 ; 1.05]). Both models are included in the publicly available code (see **Data availability** statement in the main text).

**Supplementary Material 4: means, sample sizes, and standard deviations for the continuous variables that are used after scaling in the model described in Supplementary Material 2**

| | *N* | Mean | SD |
|---|---|---|---|
| Population density * (snails/ box) | 371 (in 87 boxes) | 12.32 | 10.09 |
| ln(Shell height (mm) ) | 371 | 2.58 | 0.05 |
| ln(Shell mass (g) ) | 347 | 4.95 | 0.19 |

* Please note that scaling is done on a dataset with one observation = one individual and not one observation = one box. The mean density is thus the density experienced by the average snail, and not the density of the average box.

**Supplementary Material 5: posterior distribution of the mean value of the individual-level BLUPs (i.e. the "behavioural latent variable", $\eta_{[\mathrm{behaviour}]j}$ in the model formula Supplementary Material 2) depending on dispersal status and activity.**

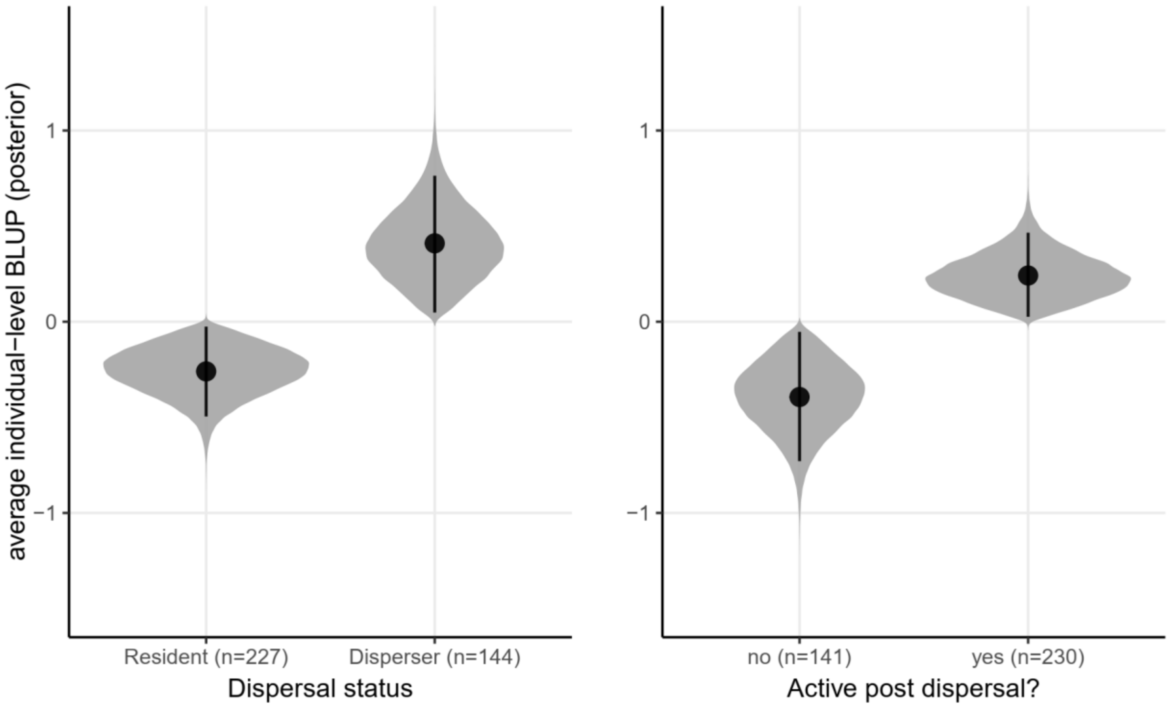

**Supplementary Material 6: pairwise posterior mean differences in relative shell mass, corresponding to the posteriors described Figure 4 in the main text.**

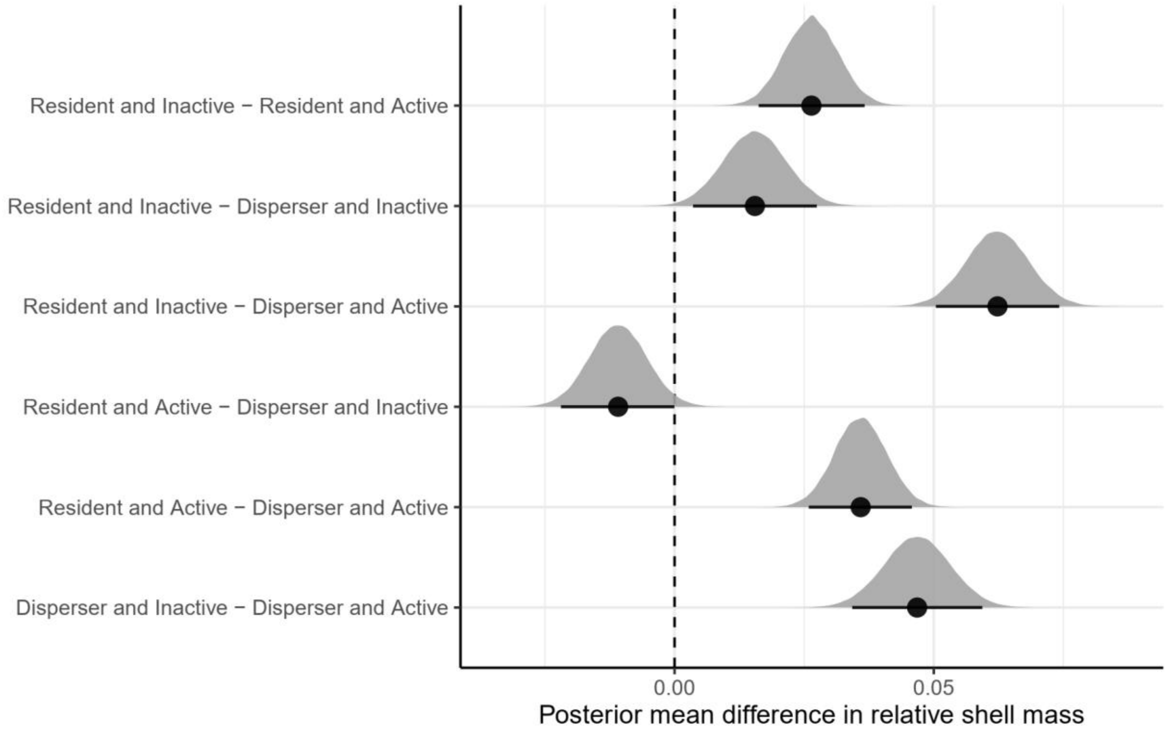


### References

Boycott, A.E. (1916). On sexual characteristics in the shell and radula of *Pomatias elegans* (Müller). *Proc. Malacol. Soc. Lond.* **v.12-13 (1916-1919)**, 127–132.

Creek, G.A. (1951). The reproductive system and embryology of the snail *Pomatias elegans* (Müller). *Proc. Zool. Soc. Lond.* **121**, 599–640.

Jordaens, K., Platts, E. & Backeljau, T. (2001). Genetic and morphological variation in the land winkle *Pomatias elegans* (Müller) (Caenogastropoda: Pomatiasidae). *J. Molluscan Stud.* **67**, 145–152.

McElreath, R. (2020). *Statistical Rethinking: A Bayesian Course with Examples in R and Stan*. 2nd edition. Boca Raton: Chapman and Hall/CRC.
